# Defense systems and prophage detection in *Streptococcus mutans* strains

**DOI:** 10.1101/2024.12.16.628709

**Authors:** O. Claisse, C. Mosterd, C. Le Marrec, J. Samot

## Abstract

Although the species is extensively studied, limited data are available on antiphage defense systems (APDSs) in *Streptococcus mutans*. The present study aimed to explore the diversity and the occurrence of APDSs and to search for prophages in the genomes of clinical isolates of *S. mutans* using bioinformatics tools.

Forty-four clinical isolates of *S. mutans* were obtained from saliva samples of people with Parkinson’s disease. Genomic DNA was extracted, sequenced using Illumina MiSeq technology, and analyzed for the presence of defense systems using DefenseFinder. CRISPR- Cas systems were characterized using CRISPRCasFinder, and prophages were detected by the PhiSpy pipeline from RAST. AcrFinder and AcrHub were used to identify anti-CRISPR proteins.

Each strain harbored between 6 and 12 APDS, with restriction-modification systems being the most prevalent, followed by the MazEF toxin-antitoxin system and CRISPR-Cas systems. Type II-C CRISPR-Cas systems were not identified here in *S. mutans*. Novel variations in type II-A signature protein Cas9 were identified, allowing their classification into four distinct groups. Variability in direct repeat sequences within the same CRISPR array was also observed, and 80% of the spacers were classified as targeting "dark matter". A unique prophage, phi_37bPJ2, was detected, showing high similarity with previously described phages. The AcrIIA5 protein encoded by phi_37bPJ2 was conserved and suggested to remain functionally active.

This study reveals the diversity of APDSs in *S. mutans* and the limited presence of prophages. The findings provide a foundation for future research on the evolutionary dynamics of these systems and their role in *S. mutans* adaptation to phage pressure.

## Introduction

*Streptococcus mutans*, a pathogen commonly associated with caries, is one of the most extensively studied Gram positive bacteria. A bibliographic search on the PubMed database in early December 2024 found over 14,800 references. However, few bacteriophages are described for this bacterium in literature. Among the bacteriophages that have been described, lytic phages predominate including e10 and f1 (for which no complete sequence is available) (Delisle and Rostkowski 1993), M102 (van der Ploeg 2007), M102AD (Delisle et al. 2012), ΦAPCM01 (Dalmasso et al. 2015) and SMHBZ8 (Ben-Zaken et al. 2021). Prophages have been described in genomes (Fu et al. 2017; Higuchi et al. 1982) but only one, ΦKSM96, has been isolated and sequenced (Sugai et al. 2023).

Despite the many articles discussing *Streptococcus mutans*, little attention has been paid to the antiphage defense systems (APDSs) of this bacterium. More data on its defense systems could help understand how this bacterium copes with the pressure exerted by bacteriophages in its environment.

In this study, we analyzed the antiphage defense systems of 44 new *S. mutans* clinical oral isolates. We also report the presence of a complete prophage whose sequence was compared with those of other *S. mutans* phages.

## Materials and Methods

### Strains: collection and identification

Clinical isolates of *Streptococcus mutans* were isolated from saliva samples of people with Parkinson’s disease. Participants were recruited in the PARKIDENT clinical trial (ClinicalTrials.gov Identifier: NCT03827551, https://clinicaltrials.gov/study/NCT03827551), whose research protocol was approved by a French regional ethics committee (approval number: Eudract N° 2018-A02773-52) and for which all participants signed a written informed consent form.

Strains were identified as previously described (Donnet et al. 2024; Ziane-Casenave et al. 2023). Briefly, saliva samples were diluted and plated on MSKB agar to isolate bacteria. Colonies were restreaked and bacteria were identified by Gram staining. *Streptococcus mutans* strains were identified by MALDI-TOF mass spectrometry.

### Genome sequencing

DNA from strains identified as *Streptococcus mutans* was extracted using the GenElute™ Bacterial Genomic DNA Kit. Complete genomic sequencing of the strains was performed on the Genome-Transcriptome platform in Bordeaux using Illumina MiSeq technology with paired reads of 2 × 300 bp. Sequences were trimmed using Trimmomatic, assembled using Skesa, and annotated through the NCBI Prokaryotic Genome Annotation Pipeline (PGAP). The complete genomes of the 44 strains used in this study are available under the Bioproject accession number: PRJNA853131.

### In silico analyses

#### Phylogenetic trees

Sequences were aligned using the Clustal Omega software (Gabler et al. 2020; Sievers and Higgins 2021). The subsequent phylogenetic trees were constructed based on the maximum likelihood method implemented in PhyML (Gabler et al. 2020; Guindon et al. 2009). Finally, the tree visualization and annotation were performed using the iTOL web tool (Letunic and Bork 2024).

#### Defense systems identification

The DefenseFinder tool (version 1.2.2) (Tesson et al. 2022) was employed to identify known antiviral defense systems among the 44 sequenced genomes.

#### CRISPR-Cas systems analysis

To identify and characterize CRISPR-Cas systems, genome sequences were analyzed using CRISPRCasFinder (Couvin et al. 2018). This tool was used to detect CRISPR arrays and classify the associated Cas genes into subtypes. Domains of Cas9 were predicted using InterPro (Blum et al. 2024) and multiple sequence alignment was performed using Clustal Omega (Gabler et al. 2020; Sievers and Higgins 2021).

#### Prophage identification

Putative prophage-like elements were identified using the PhiSpy pipeline (Akhter et al. 2012) integrated into the RAST server (Aziz et al. 2008; Brettin et al. 2015; Overbeek et al. 2014), which offers an automated platform for phage genome detection and annotation. Manual curation was subsequently performed to confirm the presence of hallmark sequences, including integrase genes, terminases, and genes encoding structural viral proteins.

#### Direct repeats and spacers in CRISPR-Cas systems

Direct repeats (DRs) and spacers within the CRISPR arrays were detected and characterized using CRISPRStudio (Dion et al. 2018) and CRISPRDetect (Biswas et al. 2016). DRs were analyzed to evaluate sequence conservation within the CRISPR-Cas system. Spacers were aligned against the Core nucleotide database using Basic Local Alignment Search Tool (BLAST) to identify potential homologs and infer their origins. Shared spacers between strains were mapped to provide an overview of the most frequently observed sequences.

#### Anti-CRISPR detection

Anti-CRISPR (Acr) elements were identified and analyzed using AcrFinder (Yi et al. 2020) and AcrHub (Wang et al. 2021). Identified Acr sequences were compared with reference Acr sequences by aligning them on the MultAlin platform (Corpet 1988).

## Results

### Defense systems in *S. mutans*

Each strain harbored between six and twelve antiphage defense systems (APDSs) (Figure 1). DefenseFinder identified twelve different APDS families in the tested strains: “Abi”, CRISPR-Cas, Dodola, Gabija, Hachiman, MazEF, a recently described family called PD (phage defense) (Vassallo et al. 2022), Retron, RloC, Restriction Modification (RM), RosmerTA, and Spbk. RM systems are the most prevalent, with types I, II, IIG, and IV detected in our genomes, sometimes all coexisting within a single strain (e.g., 12RB3, 19CLb3, and 34BRb2). Notably, all strains have at least one RM system. The MazEF toxin- antitoxin system is also present in all strains. The CRISPR-Cas and “Abi systems” follow in abundance. Three CRISPR-Cas subtypes are observed, representing both recognized classes: subtypes I-C and I-E for class 1, and subtype II-A for class 2. The “Abi systems” include members of the AbiD/F group (detected in DefenseFinder as Abi2, recently grouped together into AbiD/F; (Grafakou et al. 2024)), AbiG, AbiH, AbiJ, AbiP2, and AbiU. Apart from the APDSs mentioned above, the other systems show a variable distribution among the strains. The least represented systems are RloC (restriction-linked ORF), Hachiman and RosmerTA. The latter two are found in only two strains (07TF2 and 07TFb2), while RloC is exclusively present in strain 35DF1. Interestingly, 35DF1 is the only strain lacking an “Abi system”. Finally, the PD, Retron I-C, Dodola and Gabija systems are present in 3 to 8 strains, while the Spbk system is found in twenty strains, including the Abi-deficient strain 35DF1. Analysis of the data from these 44 strains reveals a proportion of defense systems similar to that observed for all *S. mutans* genomes sequenced to date (Figure S1).

**Figure 1:**
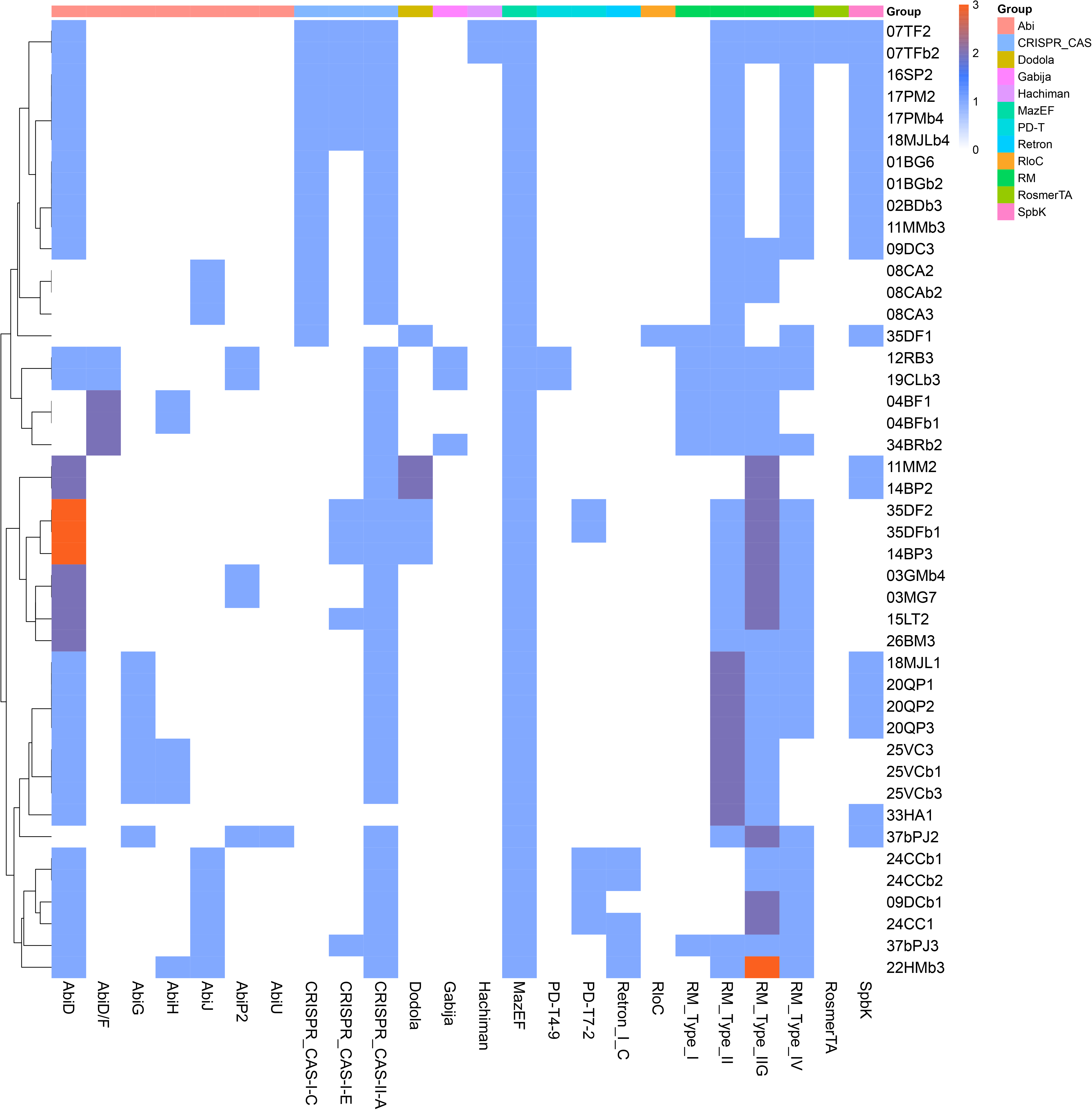
Antiphage Defense Systems identified in clinical isolates of *S. mutans* The heatmap shows the presence and frequency of antiphage defense systems in *S. mutans* strains. Each row corresponds to a bacterial strain and each column represents a specific defense system. The relative frequency of each system within a given strain is color-coded in proportion to its prevalence.

### CRISPR systems and Cas9 in *S. mutans*

Genomic analysis of the 44 strains revealed 68 CRISPR systems (Figure 2a). Almost all *Streptococcus mutans* strains (95%) possess at least the CRISPR type II-A system, with 52% of strains possessing only this system. The type I-C system is most commonly found in combination with type II-A, either alone or together with type I-E. In our study, type I-C is never found with type I-E alone, and it is the only CRISPR system present in strain 35DF1. However, type I-E is always associated with either type II-A alone or both types II-A and I-C. Notably, strain 33HA1 is unique in that it entirely lacks a CRISPR system. The presence of sequences that share high identity with cas genes in strain 37bPJ3 suggests the possible existence of a type I-A system remnant in this strain.

**Figure 2:**
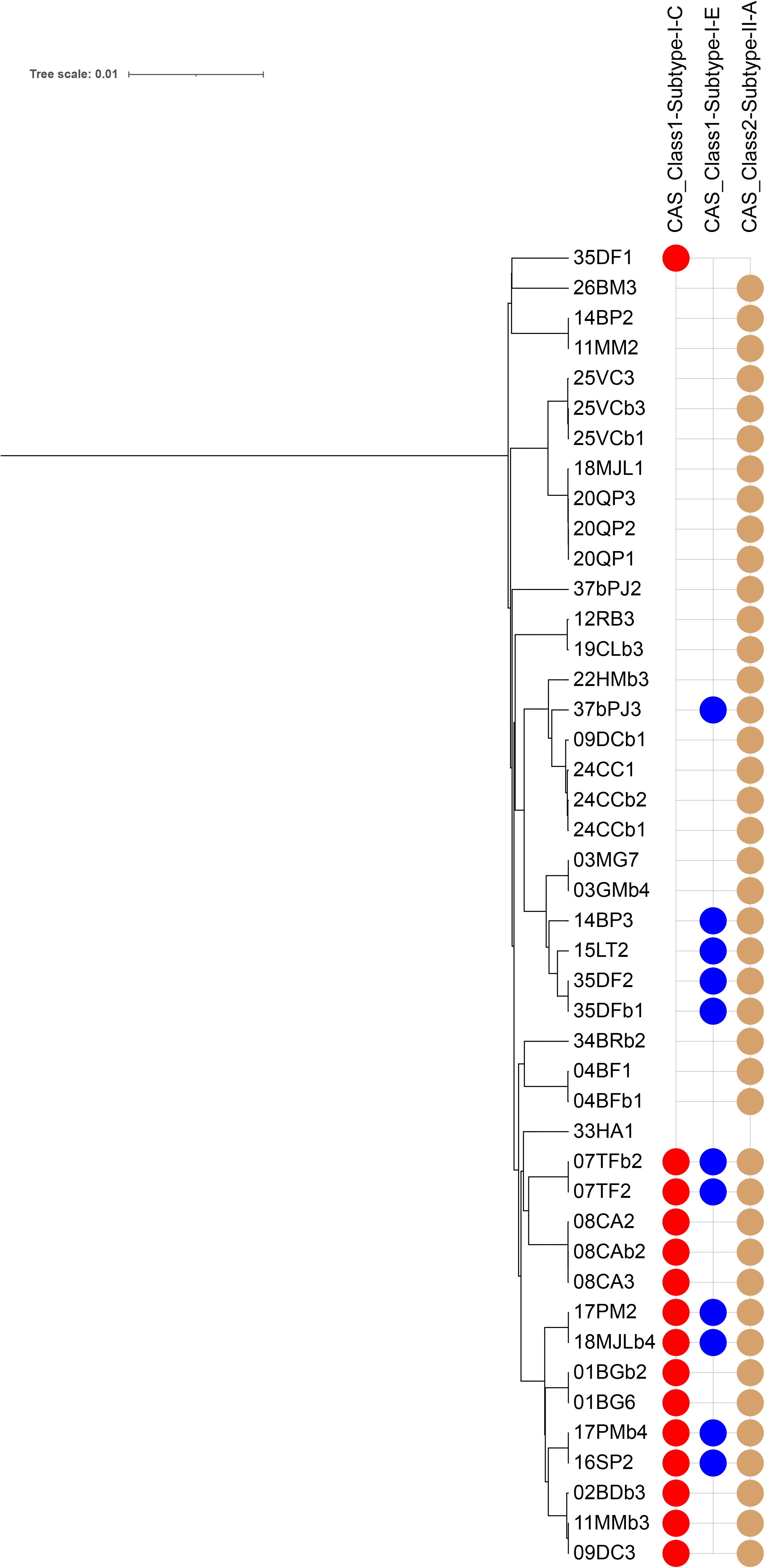
Phylogenetic Trees of CRISPR-Cas Systems and Cas9 Proteins in *S. mutans*. **(A)** Phylogenetic tree of *S. mutans* strains. Subtypes of the CRISPR-Cas systems retrieved (II- A, I-E, and I-C) are indicated at the top of the figure. The presence of one or more subtypes in a strain is represented by colored circles adjacent to the strain’s name: beige for subtype II-A, blue for subtype I-E, and red for subtype I-C. **(B)** Phylogenetic tree of Cas9 proteins from strains with type II-A CRISPR-Cas systems. Clustering highlights the sequence variations and classification of Cas9 proteins into distinct groups.

The *cas9* genes of the 42 type II-A systems were translated *in silico* and compared using Clustal Omega. This allowed for the classification of the Cas9 proteins into 4 distinct groups (figure 2b). Group 1 and Group 2 share a similar domain structure, consisting of a REC lobe, HNH endonuclease, RuvC nuclease and PAM (protospacer adjacent motif) interacting (PI) domain. As previously demonstrated, whereas the PI domains of the two groups are very different, the other domains are very similar (Mosterd and Moineau 2020).

Group 1 is the largest group (22 members, 52 % of all found Cas9 proteins) and consists of members that are all 1345 aa in size and share near identity (>98 %) over the entire protein. They are very close (>98 %) to Cas9 of the well characterized system of *S. mutans* UA159 (van der Ploeg 2007; 2009). Group 2 members (11, 26 % of all Cas9 proteins) can be further subdivided into two subgroups: 2a (7/11, 1350 aa each) and 2b (4/11, 1339 aa). The Group 2a Cas9 proteins are identical to that of the well characterized system of *S. mutans* P42S (Mosterd and Moineau 2020; 2021). Group 2a and 2b have identical PI domains. Group 2a and 2b differentiate from each other mainly by their RuvC domain (75.32 % identity when comparing 18MJL1 from 2a and 02BDb3 from 2b). The RuvC domain of Group 2b is in fact nearly identical (>98 %) to that of Group 1. The REC lobe and HNH domain are more conserved between the two subgroups (88.18 % and 90.54 % identity, respectively).

Members of Group 3 and 4 are smaller and in addition to an alpha-helical lobe, HNH and RuvC nucleases and PI domain, they possess a WED domain. Their hypothesized PI domain is smaller than that of Group 1 and 2 members. Group 3 (6 members, 14 % of all Cas9 proteins) can be further divided into two subgroups, 3a (4/6, 1125 aa) and 3b (2/6, 1126 aa), each with identical members. Group 3a and 3b share 92.25 % identity over the entire protein. When comparing only the PI domain, identity between Group 3a and 3b drops to 67.21 %. Group 4 consists of three members (7 % of all Cas9 proteins) that are identical in sequence and 1134 aa in size. Although Group 3 and 4 members share the same domain structure, they are different from each other on sequence level (67.14 % identity between Group 3a and 4, 67.03 % between 3b and 4). When comparing to Group 1 and 2, sequence identity does not exceed 21 % (Group 3) and 23 % (Group 4), respectively.

The different Cas9 proteins of *S. mutans* share similarity to Cas9 proteins from *S. thermophilus*. Group 1 and 2 members are similar to Cas9 CR3 and Group 3 and 4 members are similar to Cas9 CR1 in size, domain structure and sequence (sequence identity between CR3 and Group 1 and 2 members ranges from 61-63 % and between CR1 and Group 3 and 4 from 64-71 %).

### Prophage presence in *S. mutans*

Using the PhiSpy pipeline of RAST, we identified an intact prophage in strain 37bPJ2, which from hereon is called phi_37bPJ2 (Φ37bPJ2). The complete annotated genomic sequence of this phage is available in GenBank under accession no. JANDZM010000008.1. The sequence of phi_37bPJ2 was compared to other *S. mutans* phages, which revealed a strong resemblance (93.21 %) to prophage ctNo011 (Figure 3). Genomic regions encoding essential genes were highly conserved between the analyzed phages.

**Figure 3:**
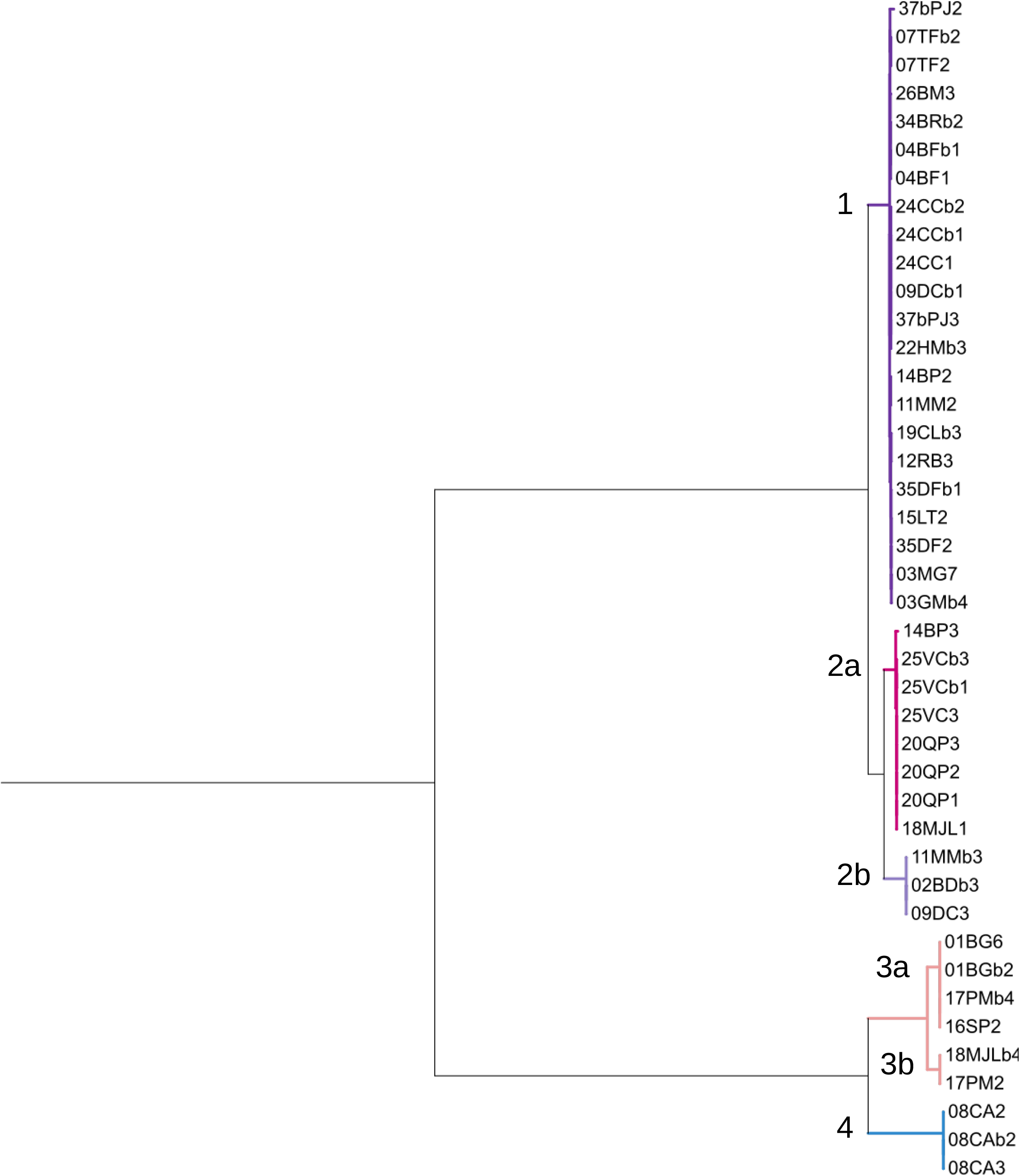
Comparison of Φ37bJ2 with other bacteriophages infecting *S. mutans*. The Φ37bJ2 sequence (JANDZM010000008.1) was compared with the sequence of the *Streptococcus mutans* temperate bacteriophage ΦKSM96 (OQ627164) isolated by Sugai et al., two previously sequenced prophages phi_MAG_isolate_ctNo011 (BK034220.1) and Phismu NLML9-1 (AHSJ01000002.1) and the virulent bacteriophage M102AD (DQ386162).

Conserved genes shared by both lytic phages and prophages exhibited sequence identities of >90%, including the tail tape measure protein (TMP) at 96–98%, small terminase at 99%, large terminase at 92–98%, endolysins at 92%, and holins at 90–92%. Notably, the integrase, a gene specific to temperate phages, showed a high level of conservation with 97% sequence identity.

### Spacers and direct repeats in CRISPR systems of *S. mutans*

Of all the DRs identified in this study, 62% are associated with CRISPR-Cas type II-A (Figure 4a). The number of different direct repeat (DR) sequences per CRISPR system varies, with a maximum of three observed. In seven strains, variations in DRs for a given CRISPR- Cas system include differences in the first four nucleotides (see Table S1 for detailed sequences). The most common configuration was two different DRs per system, representing 44% of the total. Two strains, 03MG7 and 03GMb4, possess a type II-A CRISPR system without an associated CRISPR locus. Strain 37bPJ3 appears to carry remnants of a type I-A system, with no associated CRISPR locus retrieved.

**Figure 4:**
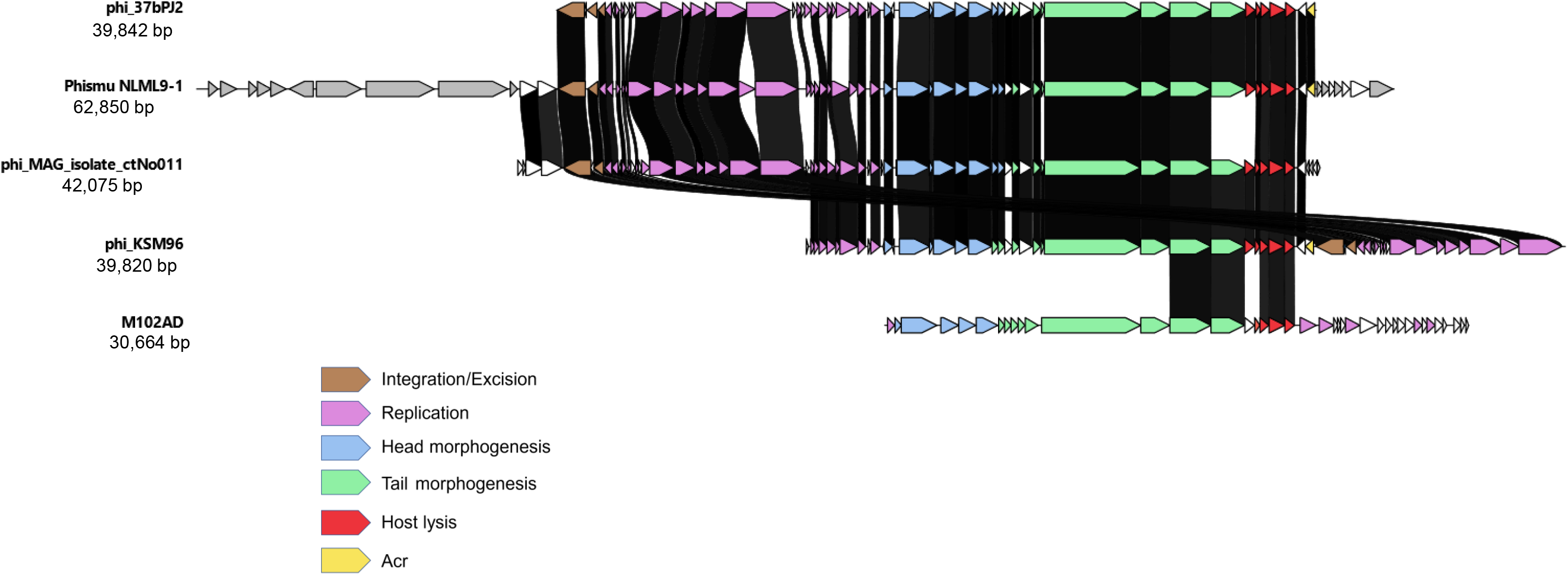
Analysis of Direct Repeat and Spacers in *S. mutans* CRISPR-Cas systems. **(A)** Stacked bar chart illustrating the distribution of direct repeat (DR) configurations within CRISPR arrays across the three described subtypes (I-E, I-C, and II-A). The bars represent the number of CRISPR arrays with 3, 2, 1, or 0 direct repeats, providing a comparative overview of DR configurations by subtype. **(B)** Network diagram showing the number of spacers identified per strain and between which strains they are shared. Shared spacers are connected by edges, while spacers with BLAST homology are highlighted in pink.

In contrast, the remaining 41 strains with an apparently complete CRISPR system contain a total of 1,059 spacers, of which 579 are unique (Figure 4b). BLAST analysis revealed homologous sequences for 117 spacers targeting different nucleotide sequences encoding terminases, TMPs or endolysins among others (see Table S2, for details). Notably, approximately 80% of the spacers matched "dark matter" sequences with no identifiable matches.

### Anti-CRISPR protein of the phage phi_37bPJ2

Through sequence analysis via AcrFinder and AcrHub, we identified an anti-CRISPR (Acr) sequence in phi_37bPJ2. The identified sequence encodes an AcrIIA5-type Acr (GenBank accession number MDT9502488.1). Sequence identity with AcrIIA5 of lytic phage D4276 targeting *S. thermophilus* is 76% (figure 5). At the N-terminus of the AcrIIA5 of phi_37bPJ2, there is a 22 amino acid segment known as the Intrinsically Disordered Region (IDR), which tends to be positively charged containing four arginine and two lysine residues. Sequence identity of this Acr with Acr from other prophages infecting *S. mutans* is over 90% (Figure S4).

**Figure 5:**
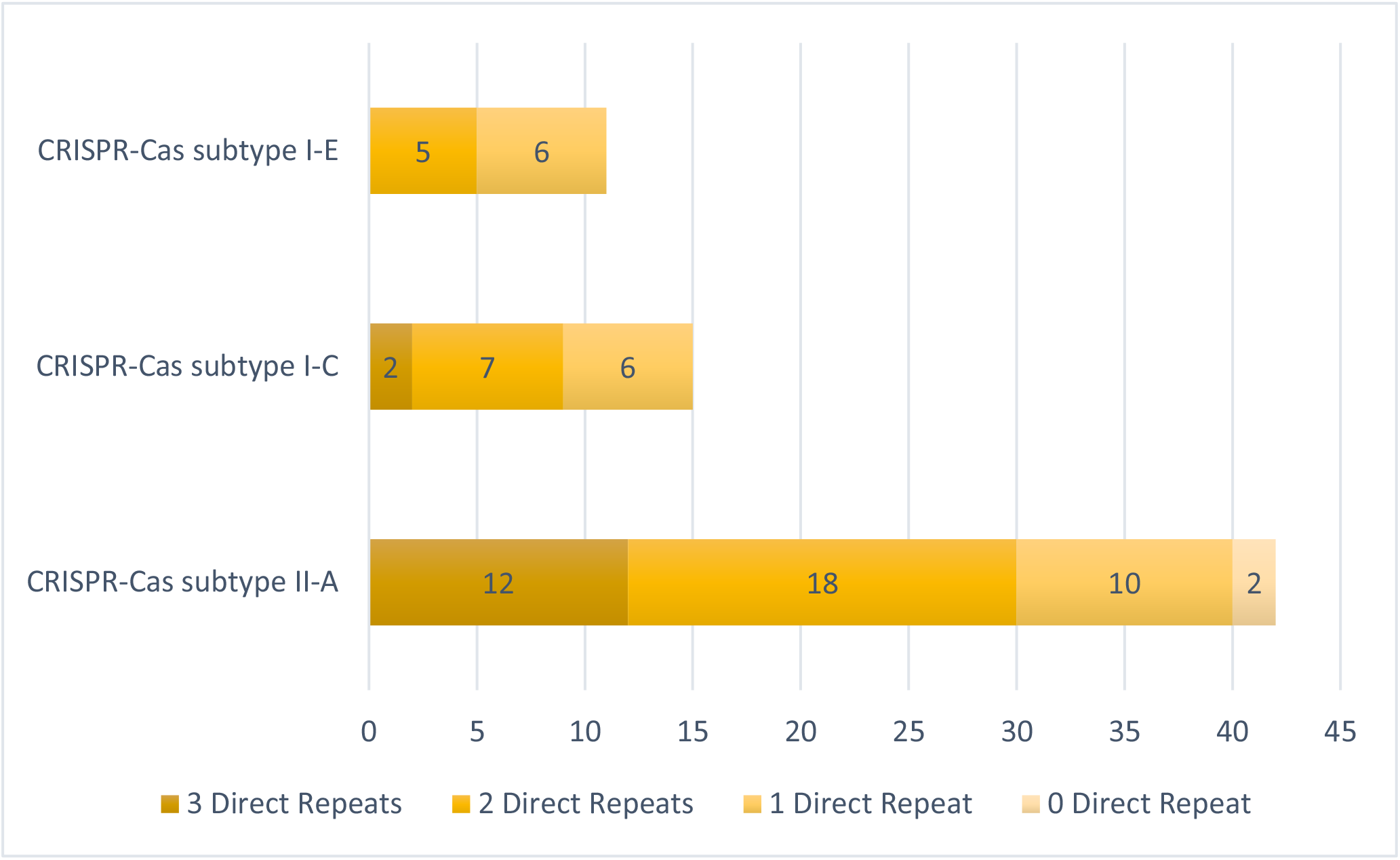
Anti-CRISPR AcrIIA5 protein identified in the phi_37bPJ2 phage. Alignment comparing the AcrIIA5 protein from prophage phi_37bPJ2 with the AcrIIA5 protein encoded by the *S. thermophilus* prophage D4276. The intrinsically disordered region (IDR) at the N-terminus, is highlighted. The alignment was performed using the Multalin tool (http://multalin.toulouse.inra.fr/multalin).

## Discussion

In this study, we identified multiple antiphage defense systems in clinical isolates of *Streptococcus mutans*, with each strain carrying between six and twelve systems. Restriction- modification (RM) systems were the most common, followed by the MazEF toxin-antitoxin and CRISPR-Cas systems. In addition, we identified a prophage in one of the strains, named phi_37bPJ2, which encodes an anti-CRISPR protein (AcrIIA5).

### Defense systems in *S. mutans*

Our study identified several antiphage defense systems in *S. mutans* but the available data on the antiphage defense systems (APDSs) of *S. mutans* remain limited.

CRISPR-Cas systems are discussed in the following paragraph. The main studies mentioning RM systems refer only to the type II RM system (Banas et al. 2011; Zhao et al. 2024). The study by Argimon et al, which also mentions RM systems, does not specify their type (Argimon et al. 2014). The RloC system has already been reported in *S. mutans*. According to Davidov et al, this system is systematically associated with the type I-C RM system (Davidov and Kaufmann 2008). In our study, the only strain with RloC was found to have a type I RM system. In this strain, we also found the Spbk system, which according to the work of Johnson et al. acts as an abortive infection system (Johnson et al. 2022). The MazEF type II toxin- antitoxin system has been previously characterized and shown to function as a growth modulator under adverse environmental conditions (Lemos et al. 2005; Syed et al. 2011).

Comparative data on other defense systems in *S. mutans* remain limited, both because of their recent discovery and the lack of comprehensive analyses across multiple strains.

However, Kelleher and colleagues recently demonstrated the presence of the Gabija, Dodola, and “Abi systems” in *Streptococcus thermophilus*, in addition to the CRISPR-Cas and RM systems, which appear to be the most important (Kelleher et al. 2024). In their study, and in contrast to our findings, the defense systems are essentially clustered in "phage defense islands". Although defense systems are commonly associated with clustering, the presence of defense islands also appeared to be absent in a study performed using *Lactococcus* species (Grafakou et al. 2024).

### CRISPR systems and Cas9 in *S. mutans*

As observed in other *Streptococcus* species relevant to human and animal health, the *Streptococcus mutans* strains in this study predominantly possess a type II-A CRISPR-Cas system (Lemaire et al. 2022). In addition, type I-C and I-E systems are also present, as previously described for *S. mutans* (Lemaire et al. 2022; Serbanescu et al. 2015; van der Ploeg 2009). In the present study, in contrast to what was reported by Lemaire et al, no type II-C was found. However, the frequency of this subtype had already been reported to be lower than the other 3 mentioned above (Lemaire et al. 2022). We also reported the absence of a CRISPR-Cas system in a strain of *S. mutans*. This has already been reported in *Streptococcus pyogenes*, but for that species, strains without CRISPR-Cas had significantly more prophages than those with it (Nozawa et al. 2011). Here, the only strain lacking CRISPR-Cas is also not lysogenic. This observation raises questions about the role of other defense systems in combating prophage integration.

In a previous study of Cas9 diversity in *S. mutans* type II-A systems, two main groups were described with conserved catalytic domains but highly variable PI domains (Mosterd and Moineau 2020). These same groups are present in our 44 strains (here named Group 1 and 2a), but several additional groups and subgroups have now been found. Group 3 and 4 consist of members that are smaller in size and with a domain structure different from what has been described in *S. mutans*. Their PI domains are different from those of characterized Cas9 proteins. Where Group 1 members recognize NGG as PAM (van der Ploeg 2009) and Group 2 members recognize NAA or NGAA (Mosterd and Moineau 2020; 2021), the PAMs targeted by Group 3 and 4 Cas9 proteins are unknown. This invites experimental validation to obtain the full picture of the PAMs recognized by *S. mutans* type II-A systems.

Most streptococcal type II-A CRISPR-Cas systems were found to be homologous to CR3 from *S. thermophilus* and CR1 homologues are rare (Lemaire et al. 2022). Previously, only CR3 homologues were described for *S. mutans* (Group 1 and 2), but with the discovery of Group 3 and 4, we demonstrate that 9 out of the 42 (21 %) type II-A systems among our *S. mutans* strains are in fact CR1 homologues. Under laboratory conditions, *S. thermophilus* CR1 is almost one tenfold more active than CR3 (Magadan et al. 2012).

### Prophage presence

So far, only five phages infecting *S. mutans* have been isolated and fully sequenced to allow complete characterization: four lytic phages (M102, M102AD, ΦAPCM01, and SMHBZ8) and one temperate phage (phi_KSM96) recently described by Sugai et al. In contrast, two additional lytic phages (e10 and f1) were reported by Delisle in 1993, but they remain unsequenced (Delisle and Rostkowski 1993). Data on prophages in *S. mutans* are similarly sparse. Fu et al. identified 35 prophages by *in silico* analysis of 171 *S. mutans* genomes, but most were incomplete, with only three having complete sequences (phismuNLML9-1, phismuN66-1, and phismu24-1) (Fu et al. 2017). Earlier, in 1977, Higuchi and colleagues reported the presence of a phage in the lysogenic *S. mutans* PK1 strain, but no genomic characterization was performed (Higuchi et al. 1977).

Genomic analysis of the strains in this study identified a unique prophage, phi_37bPJ2, which shares high sequence identity with two previously described phages: the ctNo011 prophage recovered from a human metagenome (Tisza and Buck 2021) and the phi_KSM96 phage isolated from *S. mutans* (Sugai et al. 2023). Notably, ctNo011 was initially assigned to the genus *Streptococcus* in general rather than specifically to *S. mutans*. Among the three phages, the sequence similarity is particularly pronounced in their integrase genes, suggesting that they all share the same integration site within the *S. mutans* genome. According to Sugai et al., this integration site is located between the comGB and comGC genes (or comYB and comYC, respectively, in the *S. mutans* UA159 genome) (Sugai et al. 2023). It should be noted that the contig carrying the phismuNLML9-1 phage also contains the comGC gene (data not shown).

### Spacers and direct repeat in CRISPR systems of *S. mutans*

Direct repeats (DRs) within a single CRISPR array are typically highly conserved (Jansen et al. 2002). However, variations have been observed in certain bacterial species, often limited to the final DR of the array, which is commonly described as degenerate (Biswas et al. 2016). In this study, some strains exhibited not only variations among DRs within the same CRISPR array but also differences in the first four nucleotides of the DR sequence. Such mutations in the initial nucleotides have previously been shown to affect spacer integration efficiency, potentially reducing its effectiveness (Grainy et al. 2019).

Among the spacer targets identified in this study, the majority fall into the category of "dark matter," which refers to sequences whose function or origin remains unknown because they do not match any currently known genomes or mobile genetic elements in existing databases. The proportion of "dark matter" spacers observed in our study is consistent with the results reported by Rubio et al. (Rubio et al. 2023). They introduced an innovative strategy to reduce dark matter by constructing a pangenome (a comprehensive set of genes present in all genomes of a species), which significantly reduced the proportion of unidentified spacers. Notably, most of the spacers in their analysis targeted species-specific membrane proteins. In contrast, Shmakov and colleagues primarily identified spacers targeting species-specific mobile genetic elements, including the virome (Shmakov et al. 2017; Shmakov et al. 2020). These two perspectives reflect the current main hypotheses regarding the origin of spacers: the mobilome and the host genome itself (autoimmunity).

### Anti-CRISPR (Acr) protein of the phage phi_37bPJ2

In their ongoing arms race with bacteria, phages have evolved multiple strategies to inactivate CRISPR-Cas systems, including the development of over 300 experimentally confirmed anti- CRISPR proteins (Acrs) (Wang et al. 2021). However, the efficacy of these Acrs can vary widely. For example, Garcia et al. showed that out of ten anti-CRISPR proteins tested, only AcrIIA5 effectively inhibited all type II-A and II-C Cas9 proteins tested (Garcia et al. 2019). Further research by An and colleagues suggested that AcrIIA5-mediated inhibition relies on a critical 22-amino acid segment at the N-terminus known as the Intrinsically Disordered Region (IDR). Maintaining the full length of this 22-amino acid segment is critical for its activity, as any reduction in its size significantly impairs or eliminates its ability to inhibit Cas9 (An et al. 2020). Unlike the AcrIIA5 found in the virulent phage D4276 that infects *Streptococcus thermophilus*, the IDR of the one of phi_37bPJ2 contains only six arginine or lysine residues instead of seven. These residues are critical for Acr function, and in particular the positively charged cluster of Arg12, Lys13, and Arg14 has been shown to play a key role in Cas9 inhibition (An et al. 2020). In our study, this positively charged region is well conserved in the AcrIIA5 of phi_37bPJ2. Combined with its integration into strain 37bPJ2, these findings strongly support the functional activity of AcrIIA5 in phi_37bPJ2.

The results of this study provide new insights into the diversity of antiphage defense systems in *S. mutans* and their potential role in bacterial adaptation to phage pressure. Future research should explore the functional significance of these systems, particularly CRISPR-Cas, in light of the recently identified diversity of Cas9 proteins in *S. mutans* and the frequent occurrence of AcrIIA5 in its infecting prophages. Extending genomic and functional analyses to additional clinical isolates may further elucidate the evolutionary dynamics shaping these defense systems.

## Author Contributions

OC: data curation, formal analysis, conceptualization, methodology, writing - original draft, and writing - review and editing; CM: formal analysis, conceptualization, methodology, writing - original draft, and writing - review and editing; CLM: conceptualization, writing - review and editing; JS: project administration, conceptualization, methodology, funding acquisition, investigation, formal analysis, validation, writing - original draft, and writing - review and editing

## Declaration of Conflicting Interests

The authors declare no potential conflicts of interest with respect to the research, authorship, and/or publication of this article.

## Funding

This study was supported in part by France Parkinson.

## Supplemental materials titles

**Figure S1:**

Overview of Antiphage Defense Systems identified in the genome of 478 sequenced *S. mutans* strains (May 2024 data)

Table S2: CRISPR-Cas systems and their associated direct repeats in the 44 clinical isolates of *S. mutans* from this study

Table S3: Overview of BLAST matches for 117 spacers out of 579 unique sequences: associated bacterial strains and phages

**Figure S4:**
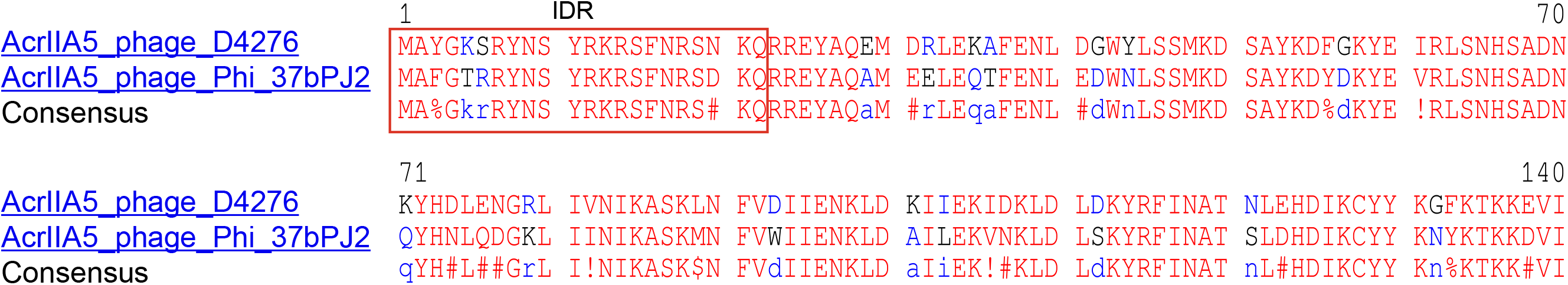
Alignment of the AcrIIA5 protein from prophage phi_37bPJ2 with AcrIIA5 proteins from other phages infecting *Streptococcus mutans*.

## Notes

### Competing Interest Statement

The authors have declared no competing interest.

## References

1. Akhter S, Aziz RK, Edwards RA. 2012. Phispy: A novel algorithm for finding prophages in bacterial genomes that combines similarity- and composition-based strategies. Nucleic Acids Res. 40(16):e126.

2. An SY, Ka D, Kim I, Kim EH, Kim NK, Bae E, Suh JY. 2020. Intrinsic disorder is essential for cas9 inhibition of anti-crispr acriia5. Nucleic Acids Res. 48(13):7584–7594.

3. Argimon S, Konganti K, Chen H, Alekseyenko AV, Brown S, Caufield PW. 2014. Comparative genomics of oral isolates of streptococcus mutans by in silico genome subtraction does not reveal accessory DNA associated with severe early childhood caries. Infect Genet Evol. 21:269–278.

4. Aziz RK, Bartels D, Best AA, DeJongh M, Disz T, Edwards RA, Formsma K, Gerdes S, Glass EM, Kubal M et al. 2008. The rast server: Rapid annotations using subsystems technology. BMC Genomics. 9:75.

5. Banas JA, Biswas S, Zhu M. 2011. Effects of DNA methylation on expression of virulence genes in streptococcus mutans. Appl Environ Microbiol. 77(20):7236–7242.

6. Ben-Zaken H, Kraitman R, Coppenhagen-Glazer S, Khalifa L, Alkalay-Oren S, Gelman D, Ben-Gal G, Beyth N, Hazan R. 2021. Isolation and characterization of streptococcus mutans phage as a possible treatment agent for caries. Viruses. 13(5).

7. Biswas A, Staals RH, Morales SE, Fineran PC, Brown CM. 2016. Crisprdetect: A flexible algorithm to define crispr arrays. BMC Genomics. 17:356.

8. Blum M, Andreeva A, Florentino LC, Chuguransky SR, Grego T, Hobbs E, Pinto BL, Orr A, Paysan-Lafosse T, Ponamareva I et al. 2024. Interpro: The protein sequence classification resource in 2025. Nucleic Acids Res.

9. Brettin T, Davis JJ, Disz T, Edwards RA, Gerdes S, Olsen GJ, Olson R, Overbeek R, Parrello B, Pusch GD et al. 2015. Rasttk: A modular and extensible implementation of the rast algorithm for building custom annotation pipelines and annotating batches of genomes. Sci Rep. 5:8365.

10. Corpet F. 1988. Multiple sequence alignment with hierarchical clustering. Nucleic Acids Res. 16(22):10881–10890.

11. Couvin D, Bernheim A, Toffano-Nioche C, Touchon M, Michalik J, Neron B, Rocha EPC, Vergnaud G, Gautheret D, Pourcel C. 2018. Crisprcasfinder, an update of crisrfinder, includes a portable version, enhanced performance and integrates search for cas proteins. Nucleic Acids Res. 46(W1):W246–W251.

12. Dalmasso M, de Haas E, Neve H, Strain R, Cousin FJ, Stockdale SR, Ross RP, Hill C. 2015. Isolation of a novel phage with activity against streptococcus mutans biofilms. PLoS One. 10(9):e0138651.

13. Davidov E, Kaufmann G. 2008. Rloc: A wobble nucleotide-excising and zinc-responsive bacterial trnase. Mol Microbiol. 69(6):1560–1574.

14. Delisle AL, Guo M, Chalmers NI, Barcak GJ, Rousseau GM, Moineau S. 2012. Biology and genome sequence of streptococcus mutans phage m102ad. Appl Environ Microbiol. 78(7):2264–2271.

15. Delisle AL, Rostkowski CA. 1993. Lytic bacteriophages of streptococcus mutans. Curr Microbiol. 27(3):163–167.

16. Dion MB, Labrie SJ, Shah SA, Moineau S. 2018. Crisprstudio: A user-friendly software for rapid crispr array visualization. Viruses. 10(11).

17. Donnet L, Claisse O, Samot J. Preprint. 2024. Serotype and distribution of adhesion genes in streptococcus mutans clinical isolates. 10.1101/2024.04.09.588668.

18. Fu T, Fan X, Long Q, Deng W, Song J, Huang E. 2017. Comparative analysis of prophages in streptococcus mutans genomes. PeerJ. 5:e4057.

19. Gabler F, Nam SZ, Till S, Mirdita M, Steinegger M, Soding J, Lupas AN, Alva V. 2020. Protein sequence analysis using the mpi bioinformatics toolkit. Curr Protoc Bioinformatics. 72(1):e108.

20. Garcia B, Lee J, Edraki A, Hidalgo-Reyes Y, Erwood S, Mir A, Trost CN, Seroussi U, Stanley SY, Cohn RD et al. 2019. Anti-crispr acriia5 potently inhibits all cas9 homologs used for genome editing. Cell Rep. 29(7):1739–1746 e1735.

21. Grafakou A, Mosterd C, Beck MH, Kelleher P, McDonnell B, de Waal PP, van Rijswijck IMH, van Peij N, Cambillau C, Mahony J et al. 2024. Discovery of antiphage systems in the lactococcal plasmidome. Nucleic Acids Res. 52(16):9760–9776.

22. Grainy J, Garrett S, Graveley BR, M PT. 2019. Crispr repeat sequences and relative spacing specify DNA integration by pyrococcus furiosus cas1 and cas2. Nucleic Acids Res. 47(14):7518–7531.

23. Guindon S, Delsuc F, Dufayard JF, Gascuel O. 2009. Estimating maximum likelihood phylogenies with phyml. Methods Mol Biol. 537:113–137.

24. Higuchi M, Higuchi M, Katayose A. 1982. Identification of pk 1 bacteriophage DNA in streptococcus mutans. J Dent Res. 61(2):439–441.

25. Higuchi M, Rhee GH, Araya S, Higuchi M. 1977. Bacteriophage deoxyribonucleic acid- induced mutation of streptococcus mutans. Infect Immun. 15(3):938–944.

26. Jansen R, Embden JD, Gaastra W, Schouls LM. 2002. Identification of genes that are associated with DNA repeats in prokaryotes. Mol Microbiol. 43(6):1565–1575.

27. Johnson CM, Harden MM, Grossman AD. 2022. Interactions between mobile genetic elements: An anti-phage gene in an integrative and conjugative element protects host cells from predation by a temperate bacteriophage. PLoS Genet. 18(2):e1010065.

28. Kelleher P, Ortiz Charneco G, Kampff Z, Diaz-Garrido N, Bottacini F, McDonnell B, Lugli GA, Ventura M, Fomenkov A, Quenee P et al. 2024. Phage defence loci of streptococcus thermophilus-tip of the anti-phage iceberg? Nucleic Acids Res.

29. Lemaire C, Le Gallou B, Lanotte P, Mereghetti L, Pastuszka A. 2022. Distribution, diversity and roles of crispr-cas systems in human and animal pathogenic streptococci. Front Microbiol. 13:828031.

30. Lemos JA, Brown TA, Jr., Abranches J, Burne RA. 2005. Characteristics of streptococcus mutans strains lacking the mazef and relbe toxin-antitoxin modules. FEMS Microbiol Lett. 253(2):251–257.

31. Letunic I, Bork P. 2024. Interactive tree of life (itol) v6: Recent updates to the phylogenetic tree display and annotation tool. Nucleic Acids Res. 52(W1):W78–W82.

32. Magadan AH, Dupuis ME, Villion M, Moineau S. 2012. Cleavage of phage DNA by the streptococcus thermophilus crispr3-cas system. PLoS One. 7(7):e40913.

33. Mosterd C, Moineau S. 2020. Characterization of a type ii-a crispr-cas system in streptococcus mutans. mSphere. 5(3).

34. Mosterd C, Moineau S. 2021. Primed crispr-cas adaptation and impaired phage adsorption in streptococcus mutans. mSphere. 6(3).

35. Nozawa T, Furukawa N, Aikawa C, Watanabe T, Haobam B, Kurokawa K, Maruyama F, Nakagawa I. 2011. Crispr inhibition of prophage acquisition in streptococcus pyogenes. PLoS One. 6(5):e19543.

36. Overbeek R, Olson R, Pusch GD, Olsen GJ, Davis JJ, Disz T, Edwards RA, Gerdes S, Parrello B, Shukla M et al. 2014. The seed and the rapid annotation of microbial genomes using subsystems technology (rast). Nucleic Acids Res. 42(Database issue):D206–214.

37. Rubio A, Sprang M, Garzon A, Moreno-Rodriguez A, Pachon-Ibanez ME, Pachon J, Andrade-Navarro MA, Perez-Pulido AJ. 2023. Analysis of bacterial pangenomes reduces crispr dark matter and reveals strong association between membranome and crispr-cas systems. Sci Adv. 9(12):eadd8911.

38. Serbanescu MA, Cordova M, Krastel K, Flick R, Beloglazova N, Latos A, Yakunin AF, Senadheera DB, Cvitkovitch DG. 2015. Role of the streptococcus mutans crispr-cas systems in immunity and cell physiology. J Bacteriol. 197(4):749–761.

39. Shmakov SA, Sitnik V, Makarova KS, Wolf YI, Severinov KV, Koonin EV. 2017. The crispr spacer space is dominated by sequences from species-specific mobilomes. mBio. 8(5).

40. Shmakov SA, Wolf YI, Savitskaya E, Severinov KV, Koonin EV. 2020. Mapping crispr spaceromes reveals vast host-specific viromes of prokaryotes. Commun Biol. 3(1):321.

41. Sievers F, Higgins DG. 2021. The clustal omega multiple alignment package. Methods Mol Biol. 2231:3–16.

42. Sugai K, Kawada-Matsuo M, Nguyen-Tra Le M, Sugawara Y, Hisatsune J, Fujiki J, Iwano H, Tanimoto K, Sugai M, Komatsuzawa H. 2023. Isolation of streptococcus mutans temperate bacteriophage with broad killing activity to s. Mutans clinical isolates. iScience. 26(12):108465.

43. Syed MA, Koyanagi S, Sharma E, Jobin MC, Yakunin AF, Levesque CM. 2011. The chromosomal mazef locus of streptococcus mutans encodes a functional type ii toxin- antitoxin addiction system. J Bacteriol. 193(5):1122–1130.

44. Tesson F, Herve A, Mordret E, Touchon M, d’Humieres C, Cury J, Bernheim A. 2022. Systematic and quantitative view of the antiviral arsenal of prokaryotes. Nat Commun. 13(1):2561.

45. Tisza MJ, Buck CB. 2021. A catalog of tens of thousands of viruses from human metagenomes reveals hidden associations with chronic diseases. Proc Natl Acad Sci U S A. 118(23).

46. van der Ploeg JR. 2007. Genome sequence of streptococcus mutans bacteriophage m102. FEMS Microbiol Lett. 275(1):130–138.

47. van der Ploeg JR. 2009. Analysis of crispr in streptococcus mutans suggests frequent occurrence of acquired immunity against infection by m102-like bacteriophages. Microbiology (Reading). 155(Pt 6):1966–1976.

48. Vassallo CN, Doering CR, Littlehale ML, Teodoro GIC, Laub MT. 2022. A functional selection reveals previously undetected anti-phage defence systems in the e. Coli pangenome. Nat Microbiol. 7(10):1568–1579.

49. Wang J, Dai W, Li J, Li Q, Xie R, Zhang Y, Stubenrauch C, Lithgow T. 2021. Acrhub: An integrative hub for investigating, predicting and mapping anti-crispr proteins. Nucleic Acids Res. 49(D1):D630–D638.

50. Yi H, Huang L, Yang B, Gomez J, Zhang H, Yin Y. 2020. Acrfinder: Genome mining anti- crispr operons in prokaryotes and their viruses. Nucleic Acids Res. 48(W1):W358–W365.

51. Zhao H, Dufour D, Zhong J, Gong SG, Roy PH, Levesque CM. 2024. Decoding adenine DNA methylation effects in streptococcus mutans: Insights into self-DNA protection and autoaggregation. Mol Oral Microbiol.

52. Ziane-Casenave S, Claisse O, Saint-Marc M, Badet M-C, Samot J. Preprint. 2023. Comparison of different methods of identification of streptococcus mutans clinical isolates. 10.2139/ssrn.4477726.

