## Supplementary figures and images for "Defense systems and prophage detection in *Streptococcus mutans* strains"

### Figure S1

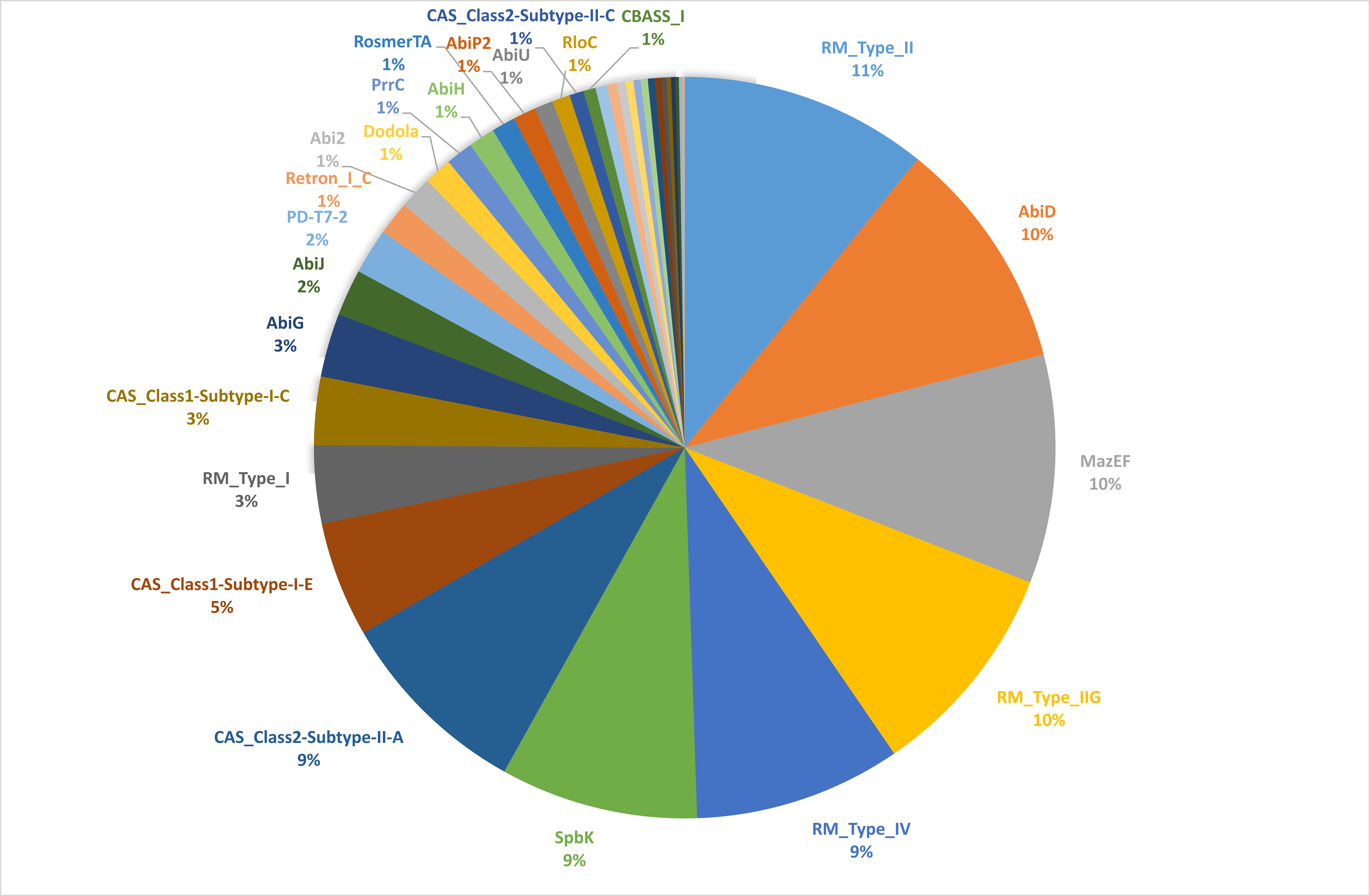
