## Supplementary material for "Defense systems and prophage detection in *Streptococcus mutans* strains": Table S2

| Strain | CAS_Type (spacer count) | Direct repeats | | | phage | Acr |
|---|---|---|---|---|---|---|
| 01BG6 | II-A (8) | gtttttgtactctcaagatttaagtaactgtacaac | | | | |
| | I-C (22) | gtcgcacccttcacgggtgcgtggattgaaat | | | | |
| 01BGb2 | II-A (8) | gtttttgtactctcaagatttaagtaactgtacaac | | | | |
| | I-C (22) | gtcgcacccttcacgggtgcgtggattgaaat | | | | |
| 02BDb3 | II-A (15) | gttttagagc**tg**tgttgtt**tcgaatgg**t**t**c**c**aaaa**c** | gttttagagc**ca**tgttagtt**actgatt**t**a**c**t**aaaa**t** | | | |
| | I-C (2) | **cc**gtcgcacccttcacgggtgcgtggattgaaata | **ac**gtcgcacccttcacgggtgcgtggattgaaata | **ag**gtcgcacccttcacgggtgcgtggattgaaat**t** | | |
| 03GMb4 | II-A (0) | | | | | |
| 03MG7 | II-A (0) | | | | | |
| 04BF1 | II-A (20) | gttttagagctgtgttgtttcgaatggttccaaaac | gttttagagctgtgttgtttcgaatggttccaaaat | gttttagagccatgttagttactgatttactaaaac | | |
| 04BFb1 | II-A (20) | gttttagagctgtgttgtttcgaatggttccaaaac | gttttagagctgtgttgtttcgaatggttccaaaat | gttttagagccatgttagttactgatttactaaaac | | |
| 07TF2 | II-A (29) | gttttagagctgtgttgtttcgaatggttccaaaac | | | | |
| | I-C (11) | gtcgcacccttcacgggtgcgtggattgaaat | | | | |
| | I-E (26) | attttacccgcacgagcgggggtgatcc | | | | |
| 07TFb2 | II-A (26) | gttttagagctgtgttgtttcgaatggttccaaaac | gttttagagctatgttgtttcgaatggttccaaaac | | | |
| | I-C (11) | gtcgcacccttcacgggtgcgtggattgaaat | | | | |
| | I-E (24) | attttacccgcacgagcgggggtgatcc | | | | |
| 08CA2 | II-A (5) | gtttttgtactctcaagatttaagtaactgtacaac | | | | |
| | I-C (12) | gtcgcacccttcacgggtgcgtggattgaaat | gtcgcaccctttaaaggttgggtttgcttttt | | | |
| 08CA3 | II-A (5) | gtttttgtactctcaagatttaagtaactgtacaac | | | | |
| | I-C (12) | gtcgcacccttcacgggtgcgtggattgaaat | gtcgcaccctttaaaggttgggtttgcttttt | | | |
| 08CAb2 | II-A (5) | gtttttgtactctcaagatttaagtaactgtacaac | | | | |
| | I-C (12) | gtcgcacccttcacgggtgcgtggattgaaat | gtcgcaccctttaaaggttgggtttgcttttt | | | |
| 09DC3 | II-A (15) | gttttagagctgtgttgtttcgaatggttccaaaac | | | | |
| | I-C (9) | gtcgcacccttcacgggtgcgtggattgaaat | | | | |
| 09DCb1 | II-A (30) | gttttagagctgtgttgtttcgaatggttccaaaac | | | | |
| 11MM2 | II-A (12) | gttttagagctgtgttgtttcgaatggttccaaaac | gttttagagccatgttagttactgatttactaaaac | | | |
| 11MMb3 | II-A (15) | gttttagagctgtgttgtttcgaatggttccaaaac | gttttggaaccattcgaaacaacacagctctaaaac | | | |
| | I-C (9) | gtcgcacccttcacgggtgcgtggattgaaat | | | | |
| 12RB3 | II-A (7) | gttttagagctgtgttgtttcgaatggttccaaaac | gttttagagccatgttagttactgatttactaaaat | | | |
| 14BP2 | II-A (12) | gttttagagctgtgttgtttcgaatggttccaaaac | gttttagagccatgttagttactgatttactaaaac | | | |

| | | | | |
|---|---|---|---|---|
| 14BP3 | II-A (2) | gttttagagctgtgttgtttcgaatggttccaaaac | gttttagagctgtgttgtttcgaatggttccaaaat | |
| | I-E (16) | attttacccgcacgagcgggggtgatcc | | |
| 15LT2 | II-A (35) | gttttagagctgtgttgtttcgaatggttccaaaac | gttttagagctgtgttgtttcgaatggttccaaaat | gttttagagccatgttagttactgatttactaaaac |
| | I-E (19) | attttacccgcacgagcgggggtgatcc | | |
| 16SP2 | II-A (10) | gttgtacagttacttaaatcttgagagtacaaaaac | gttgtacagttacttaaatcttgagagtacaaaaaa | ggatcacccccgctcgtgcgggtaaaat |
| | I-C (15) | gtcgcacccttcacgggtgcgtggattgaaat | gtcgcacccttcacgggtgcgtgggttgaaat | |
| | I-E (76) | attttacccgcacgagcgggggtgatcc | | |
| 17PM2 | II-A (29) | gtttttgtactctcaagatttaagtaactgtacaac | | |
| | I-C (27) | gtcgcacccttcacgggtgcgtggattgaaat | gtcgcaccctttaaaggttgggtttgtttttt | |
| | I-E (15) | attttacccgcacgagcgggggtgatcc | attttactcgcacgagcgggggtgatcc | |
| 17PMb4 | II-A (10) | gtttttgtactctcaagatttaagtaactgtacaac | ttttttgtactctcaagatttaagtaactgtacaac | |
| | I-C (15) | gtcgcacccttcacgggtgcgtggattgaaat | gtcgcacccttcacgggtgcgtgggttgaaat | |
| | I-E (83) | attttacccgcacgagcgggggtgatcc | | |
| 18MJL1 | II-A (8) | gttttagtaaatcagtaactaacatggctctaaaac | attttggaaccattcgaaacaacacagctctaaaac | gttttggaaccattcgaaacaacacagctctaaaac |
| 18MJLb4 | II-A (29) | gtttttgtactctcaagatttaagtaactgtacaac | | |
| | I-C (27) | gtcgcacccttcacgggtgcgtggattgaaat | gtcgcaccctttaaaggttgggtttgtttttt | |
| | I-E (15) | attttacccgcacgagcgggggtgatcc | attttactcgcacgagcgggggtgatcc | |
| 19CLb3 | II-A (8) | gttttagagctgtgttgtttcgaatggttccaaaac | gttttagagccatgttagttactgatttactaaaat | |
| 20QP1 | II-A (7) | gttttagtaaatcagtaactaacatggctctaaaac | attttggaaccattcgaaacaacacagctctaaaac | gttttggaaccattcgaaacaacacagctctaaaac |
| 20QP2 | II-A (7) | gttttagtaaatcagtaactaacatggctctaaaac | attttggaaccattcgaaacaacacagctctaaaac | gttttggaaccattcgaaacaacacagctctaaaac |
| 20QP3 | II-A (7) | gttttagagctgtgttgtttcgaatggttccaaaac | gttttagagctgtgttgtttcgaatggttccaaaat | gttttagagccatgttagttactgatttactaaaac |
| 22HMb3 | II-A (35) | gttttagagctgtgttgtttcgaatggttccaaaac | gttttagagccatgttagttactgatttactaaaac | |
| 24CC1 | II-A (18) | gttttagagctgtgttgtttcgaatggttccaaaac | gttttagagccatgttagttactgatttactaaaac | |
| 24CCb1 | II-A (23) | gttttagagctgtgttgtttcgaatggttccaaaac | gttttagagccatgttagttactgatttactaaaac | |
| 24CCb2 | II-A (24) | gttttagagctgtgttgtttcgaatggttccaaaac | gttttagagccatgttagttactgatttactaaaac | |
| 25VC3 | II-A (6) | attttggaaccattcgaaacaacacagctctaaaac | gttttggaaccattcgaaacaacacagctctaaaac | |
| 25VCb1 | II-A (6) | gttttagagctgtgttgtttcgaatggttccaaaac | gttttagagctgtgttgtttcgaatggttccaaaat | |
| 25VCb3 | II-A (6) | attttggaaccattcgaaacaacacagctctaaaac | gttttggaaccattcgaaacaacacagctctaaaac | |
| 26BM3 | II-A (11) | gttttagagctgtgttgtttcgaatggttccaaaac | gttttagagccatgttagttactgatttactaaaat | |
| 34BRb2 | II-A (5) | ggttttagagctgtgttgtttcgaatggttccaaaac | cgttttagagctgtgttgtttcgaatggttccaaaac | gttttggaaccattcgaaacaacacagctctaaaac |
| 35DF1 | I-C (2) | gtcgcacccttcacgggtgcgtggattgaaatt | gtcgcacccttcacgggtgcgtggattgaaata | gtcgcaccctttaaaggttgggtttgcttttta |
| 35DF2 | II-A (8) | gttttagagctgtgttgtttcgaatggttccaaaac | gttttagagctgtgttgtttcgaatggttccaaaat | gttttagagccatgttagttactgatttactaaaac |
| | I-E (7) | attttacccgcacgagcgggggtgatccc | attttacccgcacgagcgggggtgatcct | |

| | | | | | | |
|---|---|---|---|---|---|---|
| 35DFb1 | II-A (8) | gttttagagctgtgttgtttcgaatggttccaaaac | gttttagagctgtgttgtttcgaatggttccaaaat | gttttagagccatgttagttactgatttactaaaac | | |
| | I-E (7) | attttacccgcacgagcgggggtgatccc | attttacccgcacgagcgggggtgatcct | | | |
| 37bPJ2 | II-A (6) | gttttagagctgtgttgtttcgaatggttccaaaac | gttttagagccatgttagttactgatttactaaaat | | yes | AcrIIA5 |
| 37bPJ3 | II-A (12) | gttttagagctgtgttgtttcgaatggttccaaaac | gttttagagctgtgttgtttcgaatggttccaaagc | gttttagagccatgttagttactgatttactaaaac | | |
| | I-A (0) | | | | | |
| | I-E (11) | attttacccgcacgagcgggggtgatcc | attttacccgcacgagcggaggtgatcc | | | |
| 33HA1 | nd | | | | | |
