## Supplementary material for "Defense systems and prophage detection in *Streptococcus mutans* strains": Table S3

| product | strain/phage |
|---|---|
| AAA domain protein | isolate ctNo011 |
| AAA family ATPase | phiKSM96 |
| anti-CRISPR protein | phiKSM96 |
| ATP-binding protein | phiKSM96 |
| bifunctional DnaQ family exonuclease/ATP-dependent helicase | LAB761 |
| capsid and scaffold protein | smHBZ8 |
| capsid protein | APCM01 |
| class I SAM-dependent methyltransferase | phiKSM96 |
| Clp protease | phiKSM96 |
| conjugal transfer protein | NCH105 |
| DEAD/DEAH box helicase | KCOM 1054 |
| DEAD/DEAH box helicase | KCOM 1054 |
| distal tail protein | ctNo011 |
| DNA cytosine methyltransferase | phiKSM96 |
| DnaC-like helicase loader | M102AD |
| endolysin | M102 |
| formate C-acetyltransferase | LAB761 |
| glutamate synthase large subunit | LAB761 |
| head-tail connector protein | phiKSM96 |
| holin | APCM01 |
| hypothetical protein | KCOM 1054 |
| hypothetical protein | phiKSM96 |
| hypothetical protein | OMZ175 |
| hypothetical protein | KCOM 1054 |
| hypothetical protein | phage smHBZ8 |
| hypothetical protein | phiKSM96 |
| hypothetical protein | COCC33-14R |
| hypothetical protein | Javan237 |
| hypothetical protein | phiKSM96 |
| hypothetical protein | M102AD |
| hypothetical protein | OMZ175 |
| hypothetical protein | KCOM 1054 |
| hypothetical protein | KCOM 1054 |
| hypothetical protein | phiKSM96 |
| hypothetical protein" | UA159 |
| ImmA/IrrE family metallo-endopeptidase | KCOM 1054 |
| intergenic | LP13 |
| intergenic | GS-5 |
| intergenic | NCTC10832 |
| intergenic | smHBZ8 |
| intergenic | APCM01 |
| intergenic | MD |
| intergenic | NCTC10832 |
| intergenic | phiKSM96 |
| intergenic | NCTC10832 |
| intergenic | NCTC10832 |
| intergenic | NCTC10832 |
| intergenic | NCTC10832 |

| | |
|---|---|
| intergenic | NCTC10832 |
| intergenic | GS-5 |
| intergenic | M102 |
| intergenic | M102 |
| intergenic | M102 |
| lantibiotic ABC transporter permease | LAB761 |
| lysin | smHBZ8 |
| lysin | phiKSM96 |
| minor tail protein | phage M102 |
| minor tail protein | phage M102 |
| minor tail protein | M102 |
| minor tail protein | M102AD |
| minor tail protein | M102 |
| minor tail protein | M102AD |
| minor tail protein | M102 |
| minor tail protein | APCM01 |
| minor tail protein | M102AD |
| phage tail protein | KCOM 1054 |
| phage tail protein | KCOM 1054 |
| phage tail protein | KCOM 1054 |
| phage tail protein | KCOM 1054 |
| phage tail protein | KCOM 1054 |
| phage tail protein | KCOM 1054 |
| portal protein | phiKSM96 |
| portal protein | phiKSM96 |
| portal protein | phiKSM96 |
| predicted ATPases | LP13 |
| Protein of unknown function (DUF669) | ctNo011 |
| putative endolysin | phage M102 |
| putative major tail protein | M102 |
| putative minor structural protein | M102 |
| putative phage capsid protein | phage M102 |
| putative portal protein | M102AD |
| putative tape measure protein | M102AD |
| recombinase family protein | phiKSM96 |
| Regulatory protein repA | ctHbp13 |
| ribonuclease Y | LAB761 |
| single strand DNA binding protein | M102 |
| single-stranded DNA-binding protein | smHBZ8 |
| site-specific DNA-methyltransferase | strain S1 |
| site-specific DNA-methyltransferase | strain S1 |
| spacer | strain MD |
| spacer | strain MD |
| spacer | FDAARGOS 1458 |
| spacer | LP13 |
| spacer | LP13 |
| spacer | FDAARGOS 1458 |
| spacer | FDAARGOS 1458 |
| spacer | NN2025 |
| spacer | DPC6143 |

| | |
|---|---|
| spacer | DPC6143 |
| STRUCTURAL MAINTENANCE OF CHROMOSOMES PROTEIN | ctNo011 |
| sugar ABC transporter permease | LAB761 |
| tail family protein | phiKSM96 |
| tail length tape-measure protein | smHBZ8 |
| tail length tape-measure protein | smHBZ8 |
| tail protein | phiKSM96 |
| tail protein | ctNo011 |
| tail protein | M102 |
| tape measure protein | phiKSM96 |
| tape measure protein | phiKSM96 |
| tape measure protein | phiKSM96 |
| terminase large subunit | M102AD |
| terminase large subunit | smHBZ8 |
| terminase small subunit | M102AD |
| terminase small subunit | M102AD |
| Transporter | LP13 |
| transposon protein | UA159 |
| transposon protein | UA159 |
