## Supplementary material for "Defense systems and prophage detection in *Streptococcus mutans* strains": Figure S4

| | 1 | | | | | | 70 |
|---|---|---|---|---|---|---|---|
| AcrIIA5_phage_37bPJ2 | MAFGTRRYNS | YRKRSFNRSD | KQRREYAQAM | EELEQTFENL | EDWNLSSMKD | SAYKDYDKYE | VRLSNHSADN |
| AcrIIA5_phage_NLML9-1 | MAFGKRRYNS | YRKRSFNRSD | KQRREYAQAM | EELEQTFENL | EGWNLSSMKD | SAYKDYDKYE | VRLSNHSADN |
| AcrIIA5_phage_KSM96 | MAFGKRRYNS | YRKRSFNRSD | KQRREYAQAM | EELEQTFENL | EDWNLSSMKD | SAYKDYDKYE | VRLSNHSADN |
| Consensus | MAFGkRRYNS | YRKRSFNRSD | KQRREYAQAM | EELEQTFENL | EdWNLSSMKD | SAYKDYDKYE | VRLSNHSADN |

| | 71 | | | | | | 140 |
|---|---|---|---|---|---|---|---|
| AcrIIA5_phage_37bPJ2 | QYHNLQDGKL | IINIKASKMN | FVWIIENKLD | AILEKVNKLD | LSKYRFINAT | SLDHDIKCYY | KNYKTKKDVI |
| AcrIIA5_phage_NLML9-1 | QYHNLQDGKL | IINIKASKMN | FVWIIENKLD | AILEKVNKLD | LSKYRFINAT | SLDHDIKCYY | KNYKTKKDVI |
| AcrIIA5_phage_KSM96 | QYHNLQYGKL | IINIKASKMN | FVWIIENKLD | AILEKVNKLD | LSKYRFINAT | SLDHDIKCYY | KNYKTKKDVI |
| Consensus | QYHNLQdGKL | IINIKASKMN | FVWIIENKLD | AILEKVNKLD | LSKYRFINAT | SLDHDIKCYY | KNYKTKKDVI |
